# B cells engineered to express pathogen-specific antibodies using CRISPR/Cas9 protect against infection

**DOI:** 10.1101/541979

**Authors:** Howell F. Moffett, Carson K. Harms, Kristin S. Fitzpatrick, Marti R. Tooley, Jim Boonyaratankornkit, Justin J. Taylor

## Abstract

Effective vaccines inducing lifelong protection against many important infections such as respiratory syncytial virus (RSV), human immunodeficiency virus (HIV), influenza and Epstein-Barr virus (EBV) are not yet available despite decades of research. As an alternative to a protective vaccine we developed a genetic engineering strategy in which CRISPR/Cas9 was utilized to replace endogenously-encoded antibodies with antibodies protective against RSV, HIV, influenza or EBV in primary human or murine B cells. The engineered antibodies were expressed in up to 59% of primary B cells under the control of endogenous regulatory elements, which maintained normal antibody expression and secretion. Importantly, a single transfer of murine B cells engineered to express an antibody protective against RSV resulted in potent and durable protection against RSV infection in immunocompromised hosts. This approach offers the opportunity to achieve sterilizing immunity against pathogens for which traditional vaccination has failed to induce or maintain protective antibody responses.

---

Protective vaccines have reduced morbidity and mortality from several infectious diseases in large part by activating the humoral immune response and subsequent production of high affinity pathogen-specific antibodies produced by B cells. Unfortunately, vaccines for many common diseases are not yet available despite considerable research efforts. One example is RSV, a common pathogen that infects the upper and lower respiratory tracts. RSV is a serious threat to infants, the elderly, those with cardiopulmonary disease, and those undergoing hematopoietic stem cell transplant, where it is a significant cause of morbidity and mortality^1–3^. The first RSV vaccine trial occurred in 1966 but proved harmful when subsequent natural infection caused severe lower respiratory disease and two deaths^4^. The second RSV vaccine trial occurred in 2016 and also failed to protect against infection^5^. While several other candidate RSV vaccines are being developed, it is unclear if any will elicit a protective response.

Despite the failure of vaccine trials, evidence exists for antibody-mediated protection against RSV. Several monoclonal antibodies including the RSV-specific monoclonal antibody palivizumab have been shown to protect against RSV infection *in vitro* or *in vivo*^6,7^. Similarly, monoclonal antibodies protective against HIV, influenza, EBV, human metapneumovirus virus, Dengue, Zika, Ebola and many other pathogens are also being developed^8^. However, the infusion of monoclonal antibodies like palivizumab is limited to high risk populations because monthly reinfusion is required to maintain protection. While new approaches to increase the antibody half-life after injection have been developed^9^, even the most promising of these strategies would require lifelong reinfusion to maintain protection.

To overcome the need for reinfusion, alternative strategies to generate long-term immunity have been explored. One approach involves viral transduction of muscle cells with an adenoviral vector encoding a protective antibody^10,11^. Another approach is transduction of hematopoietic stem cells with a lentivirus-encoded secreted antibody, which are differentiated into antibody-secreting plasma cells *in vitro* prior to infusion, or allowed to differentiate *in vivo* after infusion^12,13^. A shared limitation to both the adenoviral/muscle and lentiviral/stem cell approaches is that the level of antibody produced is fixed and unresponsive to infection. In contrast, protective vaccines elicit both long-lived memory B cells and antibody-secreting plasma cells. Memory B cells express a membrane bound form of antibody that allows these cells rapidly respond and differentiate into additional antibody-secreting cells upon infection.

In an effort to mimic the protective B cell response, we developed a genetic engineering strategy that allowed for the expression of protective antibodies against RSV, HIV, influenza or EBV in mouse or human B cells under endogenous regulatory elements. This was challenging because fully functional B cells require alternative splicing and polyadenylation to produce membrane bound as well as secreted antibody, a process which is difficult to recapitulate in a viral transgene^14,15^. Adding an additional level of difficulty, antibodies are produced as the product of two genes, heavy chain gene (*IgH*) and either the kappa (*IgL-κ*) or lambda (*IgL-λ*) light chain gene. Targeting the heavy chain locus was complicated by the large size and extreme genetic heterogeneity of this area in antibody-expressing B cells. Each developing B cell undergoes recombination of *V*, *D*, and *J* segments over more than a megabase of DNA within the heavy chain locus, and this results in variable regions that are essentially unique to each cell^16^. This sequence variability makes directly targeting antibody coding regions challenging, and replacement of the entire heavy chain locus has been inefficient to date^17^. To bypass this complexity, we developed a single cut approach where the full light chain linked to the heavy chain VDJ was inserted into an intronic region of the heavy chain locus. Using this approach we find that both murine and human B cells can be efficiently engineered and a single transfer of murine B cells engineered to express an RSV-specific antibody can protect a recipient animal from infection for several months.

## Results

### Targeting strategy and emAb cassette design

To circumvent the complexity of the antibody heavy chain gene, we focused upon a small 2600 nucleotide region of DNA present in all B cells between the last *J* gene segment and the region involved in class switching. This region was further limited due to the presence of a critical intronic Eμ enhancer, one of several strong enhancer elements that cooperates to drive high level expression of recombined *VDJ* genes despite the weak core promoters of V gene segments^18,19^. Activity of these enhancers is regulated in part by the proximity of promoters relative to the Eμ enhancer, and insertion of a transgene between the recombined VDJ segments and the Eμ enhancer can completely block transcription of the upstream VDJ segment^20^. We therefore inserted a synthetic *VDJ* under the control of a heavy chain promoter upstream of the Eμ enhancer would allow for physiological expression of the inserted engineered monoclonal antibody, which we termed an “emAb”.

To enable one-hit insertion we designed an emAb cassette that contained a heavy chain promoter followed by the complete light chain linked to a recombined heavy chain VDJ containing a splice junction to allow for splicing to downstream endogenous heavy chain constant regions (Fig. 1a). Utilizing the endogenous heavy chain constant region reduced the insert size and allowed emAbs to be expressed in membrane-bound and secreted forms of all isotype classes under the control of endogenous regulatory elements. When expressed, the emAb light chain is physically linked to the heavy chain with a 57 amino acid glycine-serine linker (Fig. 1b), which has been used previously in single chain Fab fragments^21^. The linker also contained three tandem Streptag-II motifs to facilitate the detection and enrichment of engineered cells^22^. Physically linking the heavy and light chains also minimized the possibility of mispairing between an inserted emAb heavy chain and the endogenous light chain.

**Figure 1.**
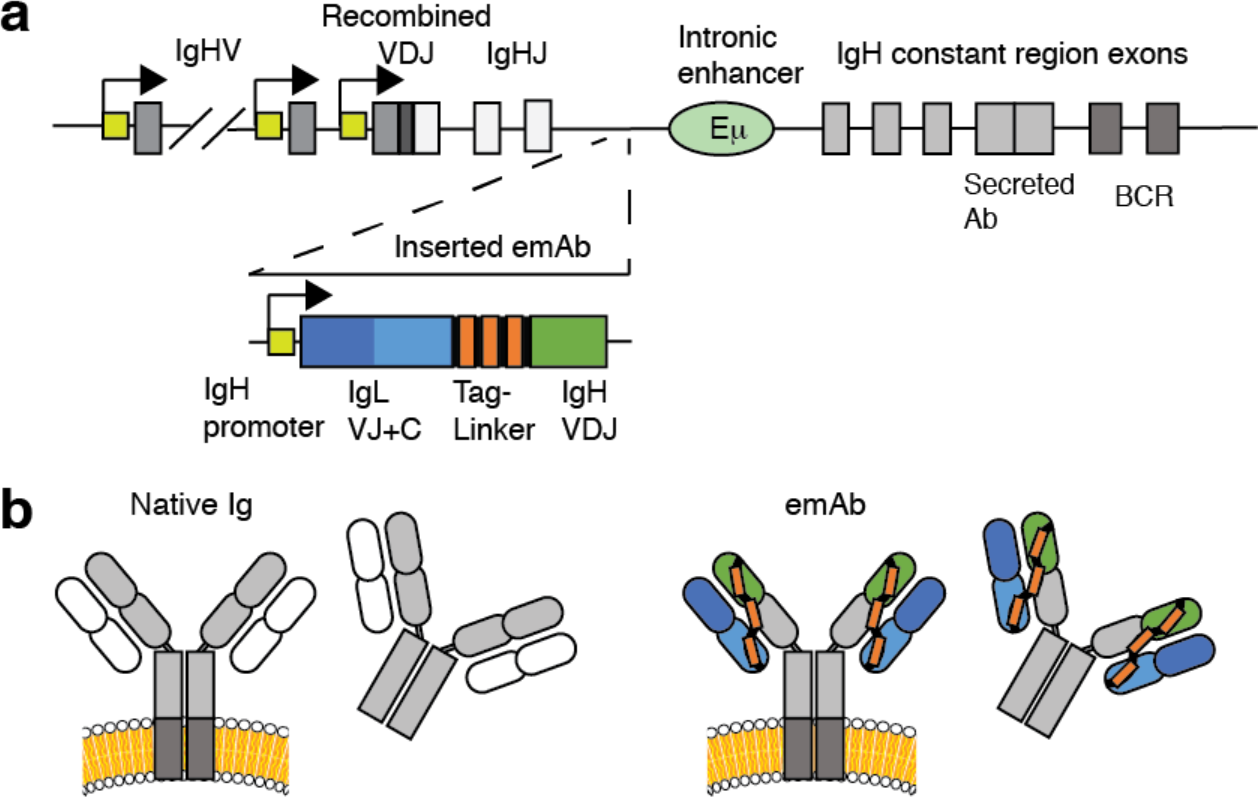
CRISPR/Cas9 mediated replacement of an endogenous antibody with an engineered monoclonal antibody (emAb). **(a)** Diagram showing insertion of an emAb cassette containing a heavy chain promoter, full light chain, linker, and partial heavy chain into the intronic region between the terminal J segment and the Eμ enhancer, upstream of the endogenous heavy chain constant region exons. **(b)** Schematic showing normal transmembrane and secreted antibody versus transmembrane and secreted emAb with the endogenous heavy chain constant regions shown in grey.

### CRISPR/Cas9-mediated emAb expression in RAMOS B cells

Insertion and expression of an emAb cassette was first tested in the Burkitt-lymphoma derived RAMOS B cell line that natively expresses membrane-bound and secreted antibodies. Analyzing the region between the terminal J segment and Eμ using the CrispRGold algorithm^23^, several potential Cas9 guide RNA (gRNA) binding sites were identified. We focused upon gRNA huIgH296, which targeted a region was 296 nucleotides downstream of IgHJ6 where no single nucleotide polymorphism (SNP) with a frequency above 1% have been reported^24^. Electroporation of RAMOS cells with huIgH_296_ gRNA precomplexed with Cas9 protein resulted in efficient DNA cutting, with insertions or deletions (indels) at this site being detected in ~72% of genomic heavy chain DNA (Fig. 2a). After electroporation, we incubated cells with AAV encoding an engineered αRSV-emAb cassette derived from palivizumab flanked by 450 nucleotide homology arms on either side of the huIgH_296_ target site. Since the AAV does not include the heavy chain constant regions which are essential for antibody expression, RAMOS cells would only gain the ability to bind RSV F antigen if the αRSV-emAb cassette was successfully inserted into the heavy chain locus. Flow cytometry was used to assess αRSV-emAb expression on the cell surface by measuring binding to fluorescent RSV F antigen and streptactin, a modified streptavidin with high affinity for the Streptag-II motifs in the linker^25^. Using this approach, ~30% of αRSV-emAb-engineered RAMOS cells bound RSV F antigen and streptactin compared to less than 0.3% of control RAMOS cells (Fig. 2b, c). In order to assess the functionality of the αRSV-emAb, RAMOS cells binding RSV antigen and streptactin were FACS-purified to create a αRSV-emAb^+^ cell line. To confirm functional interaction between RSV-emAb^+^ and the BCR signaling complex, control and αRSV-emAb^+^ RAMOS cells were stimulated with tetramerized RSV F antigen or with polyclonal αIg F(ab’)_2_ antibody. Only RSV-emAb^+^ cells fluxed calcium in response to RSV F antigen, whereas and αRSV-emAb^+^ and control RAMOS cells had similar responses to αIg (Fig. 2d, e). These results indicate that emAb engineering reprograms B cells with a functional monoclonal antibody.

**Figure 2.**
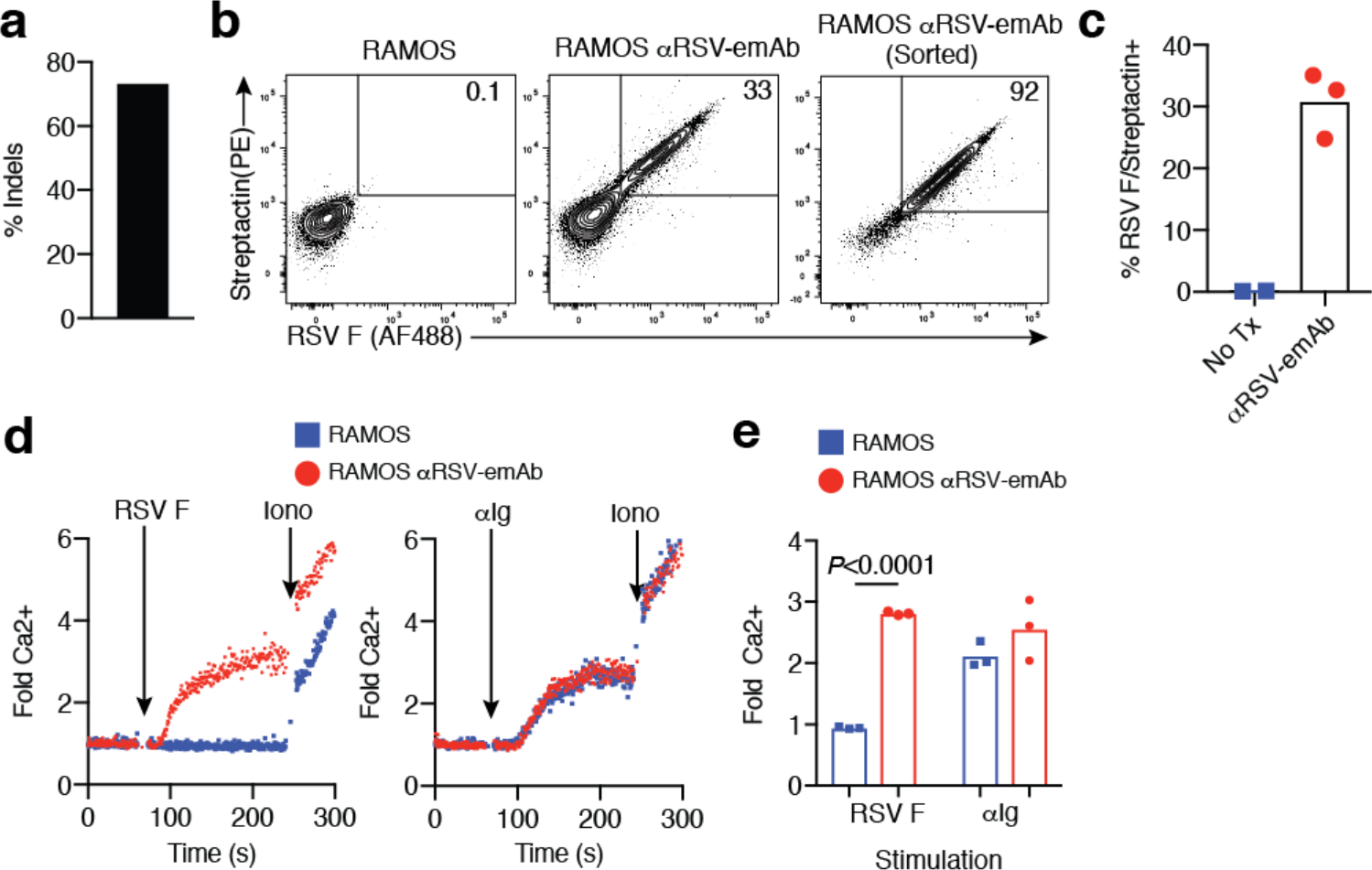
CRISPR/Cas9 mediated replacement of an endogenous antibody with an emAb targeting RSV in RAMOS B cells. **(a)** The frequency of single nucleotide polymorphisms at the huIgH_296_ gRNA cutting site analyzed by tracking of indels by decomposition. **(b)** Representative flow cytometric analysis of RSV F antigen and Streptactin binding by control RAMOS cells and engineered RAMOS cells 2 days following electroporation with huIgH_296_ gRNA complexed with Cas9 followed by incubation with an AAV encoding an αRSV-emAb cassette. Cell binding before (middle panel) and after FACS-purification and expansion (right panel) is displayed. **(c)** Percent of RAMOS cells binding both RSV F antigen and Streptactin following emAb engineering from three independent experiments. **(d)** Representative flow cytometric analysis and **(e)** peak fold increase in Fluo-4 fluorescence in αRSV-emAb^+^ RAMOS cells and control RAMOS cells following stimulation with 1 μg/mL αIg F(ab’)_2_ or 1 μg/mL tetramerized RSV F antigen. Data points are pooled from 3 independent experiments using RAMOS and sorted RAMOS RSV-emAb cell lines.

### CRISPR/Cas9-mediated emAb expression in primary human B cells

We next engineered human primary B cells using a multistep process of expansion and differentiation (Fig. 3a). Human B cells were MACS-purified from PBMCs and stimulated with a cocktail of cytokines, a multimerized CD40 ligand and CpG for 48 hours prior to electroporation with huIgH296 gRNA/Cas9. Sequencing of genomic DNA revealed the presence of indels in ~67% of heavy chain DNA sequences from six independent donors (Fig. 3b). To test the performance of the emAb backbone across multiple independently derived antibodies, we designed three additional emAb cassettes encoding for the HIV-1 broadly neutralizing antibody VRC01, the influenza broadly neutralizing antibody MEDI8852, and the EBV neutralizing antibody AMM01^26–28^. All emAb cassettes were efficiently introduced into primary human B cells and resulted in the generation of populations which bound RSV F antigen, influenza hemagglutinin (HA) stem, EBV gH/gL or HIV-1 Envelope (Env) (Fig. 3c). Tests of αRSV-emAb engineering in seven independent donors resulted in RSV antigen being bound by 16-44% of engineered B cells, compared to under 1% of mock engineered control B cells. The emAb expression efficiency ranged from 5-59% for the three additional antiviral emAbs (Fig. 3d). All four emAb B cell populations also secreted engineered antibodies when induced to differentiate into antibody-secreting cells through additional culture (Fig. 3e). These data demonstrate the flexible nature of the emAb platform for engineering primary B cells to produce and secrete protective monoclonal antibodies.

**Figure 3.**
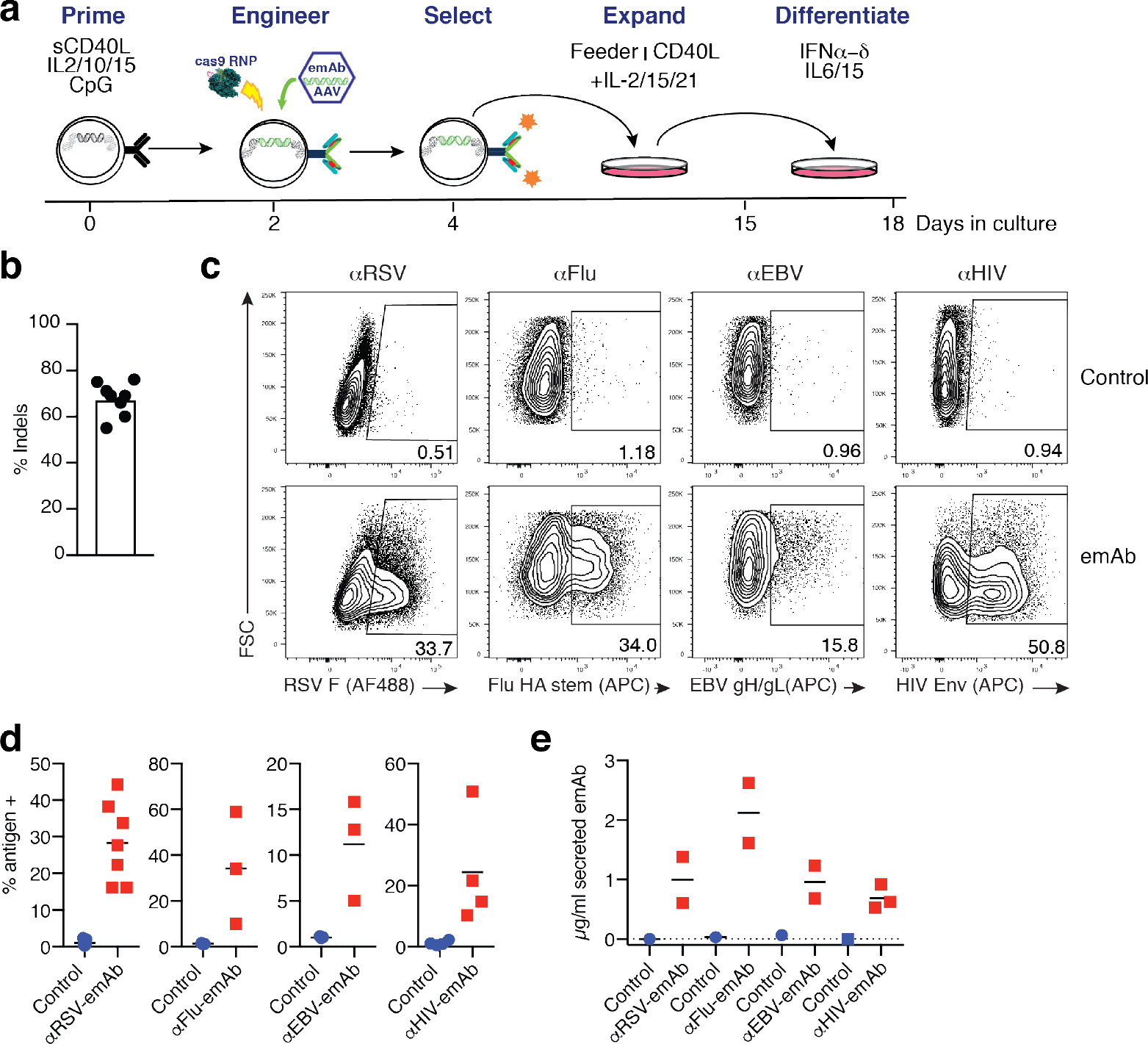
CRISPR/Cas9 mediated replacement of an endogenous antibodies with an engineered emAbs targeting RSV, HIV-1, Influenza or EBV in primary human B cells. **(a)** Schematic representation of human B cell culture and engineering. **(b)** Frequency of Indels detected in genomic heavy chain sequences from B cells electroporated with gRNA huIgH_296_/Cas9 two days prior to analysis. Each data point represents B cells from a different individual. **(c)** Representative flow cytometric analysis and **(d)** quantitation of antigen binding to human B cells engineered to express emAbs based upon the HIV-1 broadly neutralizing antibody VRC01 (αHIV-emAb), the influenza broadly neutralizing antibody MEDI8852 (αFLU-emAb), the EBV neutralizing antibody AMM01(αEBV-emAb), or αRSV-emAb. Cells were analyzed at the day four selection step shown in panel a and control B cells were mock electroporated and expanded similarly to the engineered cells. Each data point in d represents the frequency of antigen-binding B cells from a different individual (n=3-7). **(e)** ELISA-mediated quantitation of antigen-specific antibodies in day 18 supernatants from 2-3 independent B cell cultures per emAb B cell population.

### Engineered B cells can co-express emAbs off of both heavy chain loci

Work with transgenic mice demonstrated that productive VDJ sequences on both heavy chain loci results in simultaneously transcription and translation of both heavy chains^29^. These results indicate that if the emAb cassette was only inserted into the heavy chain locus that did not undergo VDJ recombination or was otherwise recombined in a way that did not produce a productive heavy chain, cells simultaneously expressing their endogenous antibodies and emAbs could be produced (Fig. 4a). This would be problematic if the endogenous antibody caused autoimmune tissue destruction due to binding self-antigens. This is a concern given that up to 20% of the naive B cell repertoire has been shown to express antibodies that can bind self-antigens^30^. To determine whether B cells co-expressing both the emAb and endogenous antibodies were produced, we FACS-purified B cells that expressed antibodies utilizing lambda light chains (Fig. 4b) prior to engineering with αFlu-emAb, which utilized a kappa light chain. Since many cells in the culture downregulated surface BCR expression as a result of the culture conditions, we gated on cells expressing high levels of CD79b to focus on cells retaining high surface BCR expression (Fig. 4b). Within the αFlu-emAb engineered CD79b^+^ B cells, many cells that gained the ability to bind Flu hemagglutinin (HA) stem antigen lost surface expression of the lambda light chain (Fig. 4b). The loss of lambda light chain expression indicated that the inserted emAb cassette blocked expression of the endogenous antibody in these cells. However, around half of Flu HA stem-binding emAb B cells retained lambda light chain expression on the cell surface (Fig. 4b, c). These results suggested that in around half of the cells, emAb insertion occurred on the non-productive heavy chain locus, resulting in co-expression of the emAb and the endogenous antibody.

**Figure 4.**
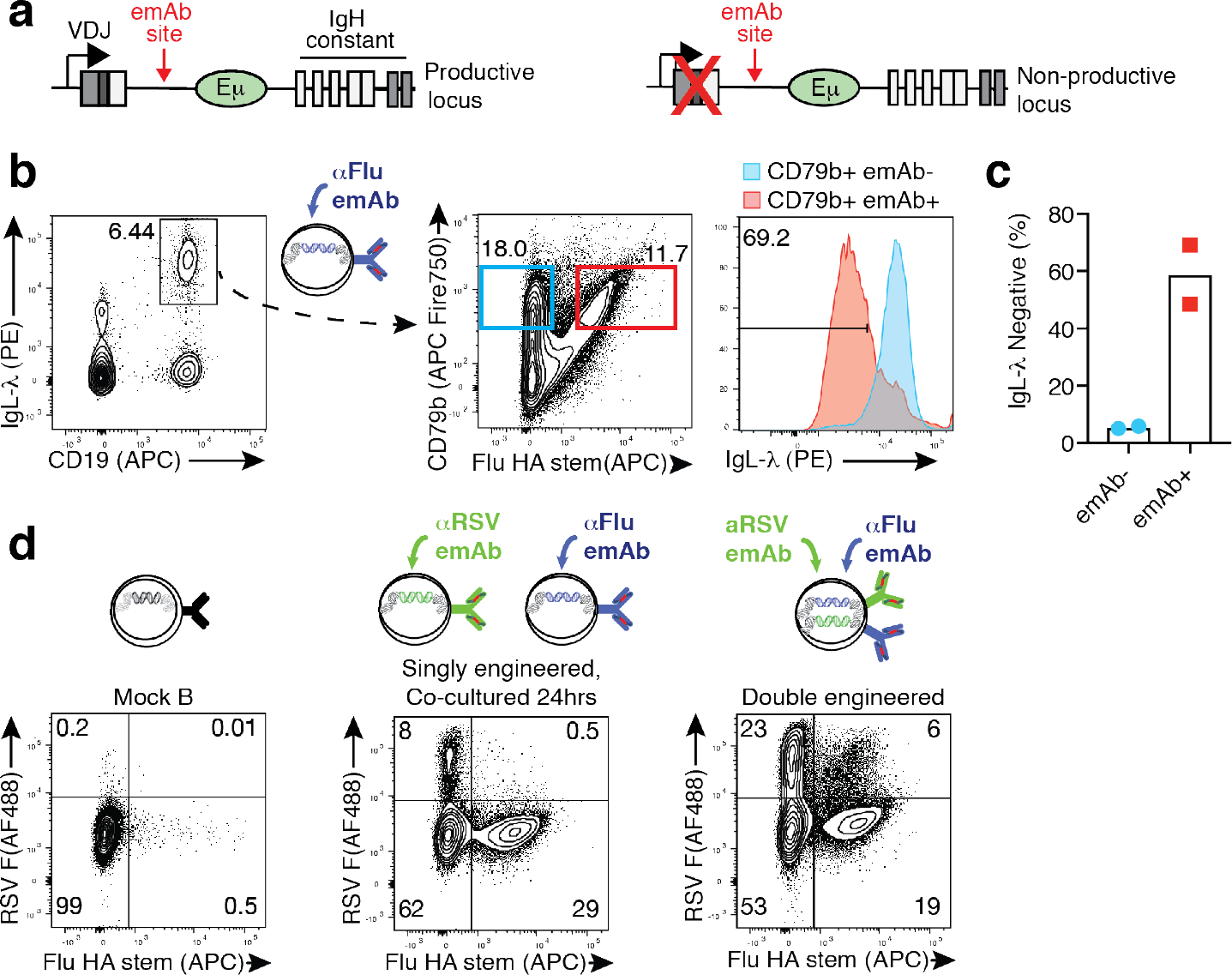
Simultaneous emAb expression from both heavy chain loci. **(a)** Diagram of a productively recombined heavy chain and a non-productive heavy chain, both of which contain the huIgH_296_ gRNA target site. **(b)** Representative flow cytometric analysis and **(c)** quantitation of the loss of IgL-λ expression by FACS-purified IgL-λ^+^ primary cells engineered to express αFlu-emAb, which utilizes IgL-κ. **(d)** Representative flow cytometric analysis of RSV F antigen and Flu HA stem antigen binding by mock engineered control B cells, B cells engineered separately with αFlu-emAb and αRSV-emAb and co-cultured for 24 hours prior to analysis, or B cells simultaneously double-engineered with αFlu-emAb and αRSV-emAb. Data are representative of two independent experiments.

The expression of emAb from both the productive and non-productive heavy chain offered the possibility of producing dual-emAb B cells by insertion of a different cassette into each locus. To test this possibility, we assessed B cells engineered simultaneously with AAVs encoding αRSV-emAb and αFlu-emAb cassettes. Simultaneous engineering of cells with αRSV-emAb and αFlu-emAb emAb resulted in ~6% of cells binding both RSV F and Flu-HA stem, a population which is not detected in control B cells, or cells that were engineered with the individual AAVs followed by co-culture (Fig. 4d). Taken together, these results demonstrated that emAbs can be simultaneously expressed by both heavy chain loci.

### Murine emAb B cells protect against RSV infection

Having demonstrated the ability to engineer primary B cells, we next assessed the protective capability of these cells in a murine model of infection. Murine emAb-expressing B cells were produced using a process of priming, electroporation and emAb cassette delivery, and expansion similar to that used in human primary B cells (Fig. 5a). Electroporation in combination with pre-complexed muIgH_367_ gRNA and Cas9 resulted in indels in ~80% of target alleles (Fig. 5b). Delivery of a murine αRSV-emAb cassette encoded by AAV resulted in 8-24% of murine B cells binding RSV antigen two days later (Fig. 5c, d). RSV antigen-binding of 1-7% of B cells could be detected when an αRSV-emAb cassette was delivered during the electroporation as a double stranded DNA (dsDNA) containing short 36 nucleotide homology regions (Fig. 5c, d), offering a potential for producing emAb B cells without the use of an AAV.

**Figure 5.**
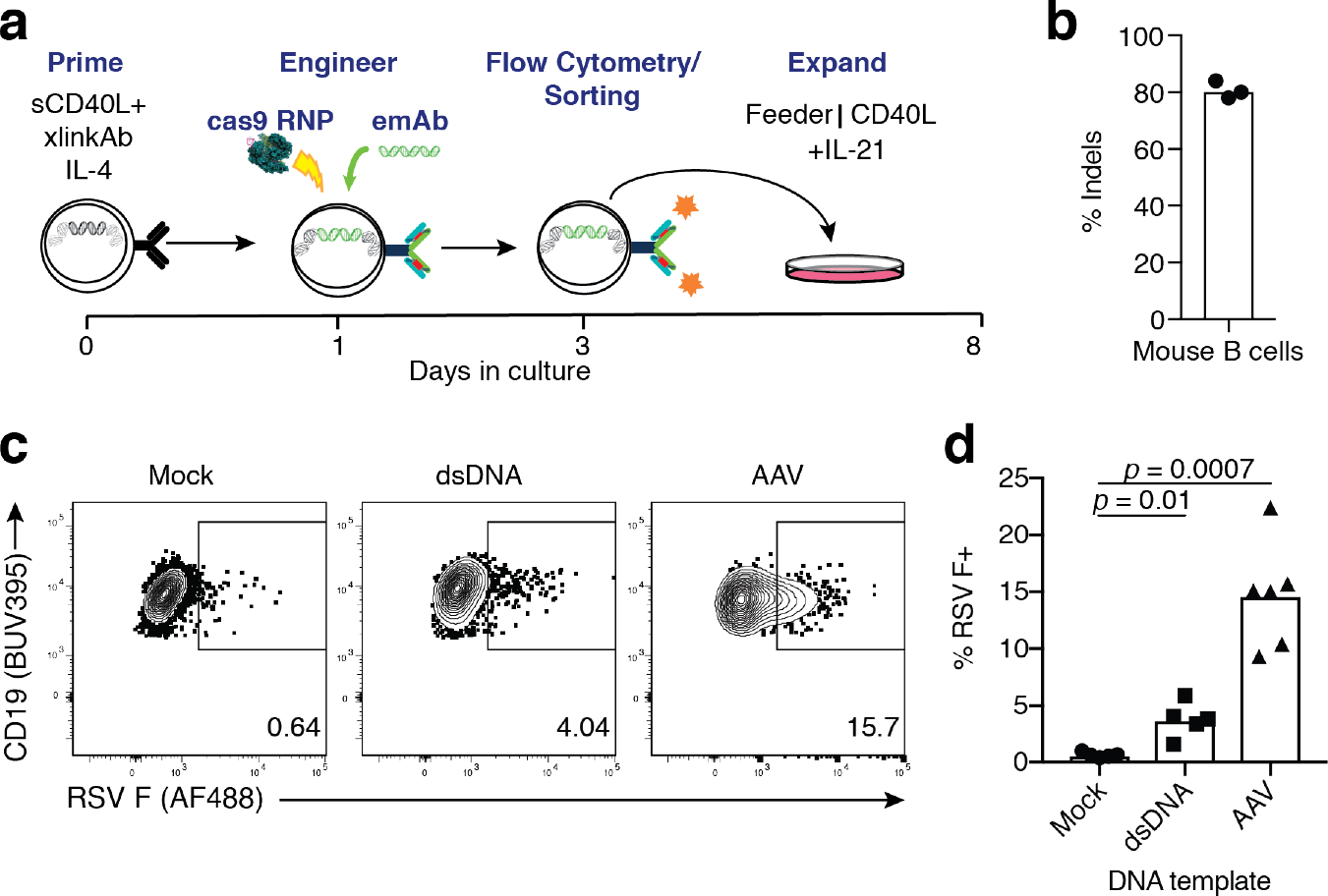
Engineering of primary mouse B cells to express αRSV-emAb. **(a)** Schematic representation of the mouse B cell culture and engineering. **(b)** Frequency of indels detected in genomic heavy chain sequences from B cells two days after electroporation with muIgH_367_ gRNA/Cas9. Each data point represents B cells from an independent experiment (n=3). **(c)** Representative flow cytometric analysis and **(d)** quantitation of RSV F antigen binding to mouse B cells engineered to express αRSV-emAb using cassettes delivered purified dsDNA or AAV. Each data point in **d** represents the frequency of antigen-binding B cells from an independent experiment (n=5-6).

To assess the protective ability of αRSV-emAb B cells, 1.5 × 10^7^ CD45.1^+^ Balb/cByJ mouse B cells binding RSV antigen were transferred into wild-type CD45.2^+^ Balb/cByJ mice (Fig. 6a). Six days following transfer, 3-9 μg/mL of RSV-specific antibodies were present in the blood of mice that received αRSV-emAb B cells, but not control B cells (Fig. 6b). Animals were subsequently challenged intranasally with 10^6^ PFU of RSV and virus was measured in the lung five days later. Around 5000 PFU of RSV could be detected in lungs from control mice that did not receive cells, and recipients of control B cells (Fig. 6c). In contrast, RSV was nearly undetectable in mice that received αRSV-emAb B cells (Fig. 6c). This protection was comparable to the protection afforded by the injection of a clinical dose, 15 mg/kg, of palivizumab two days before infection.

**Figure 6.**
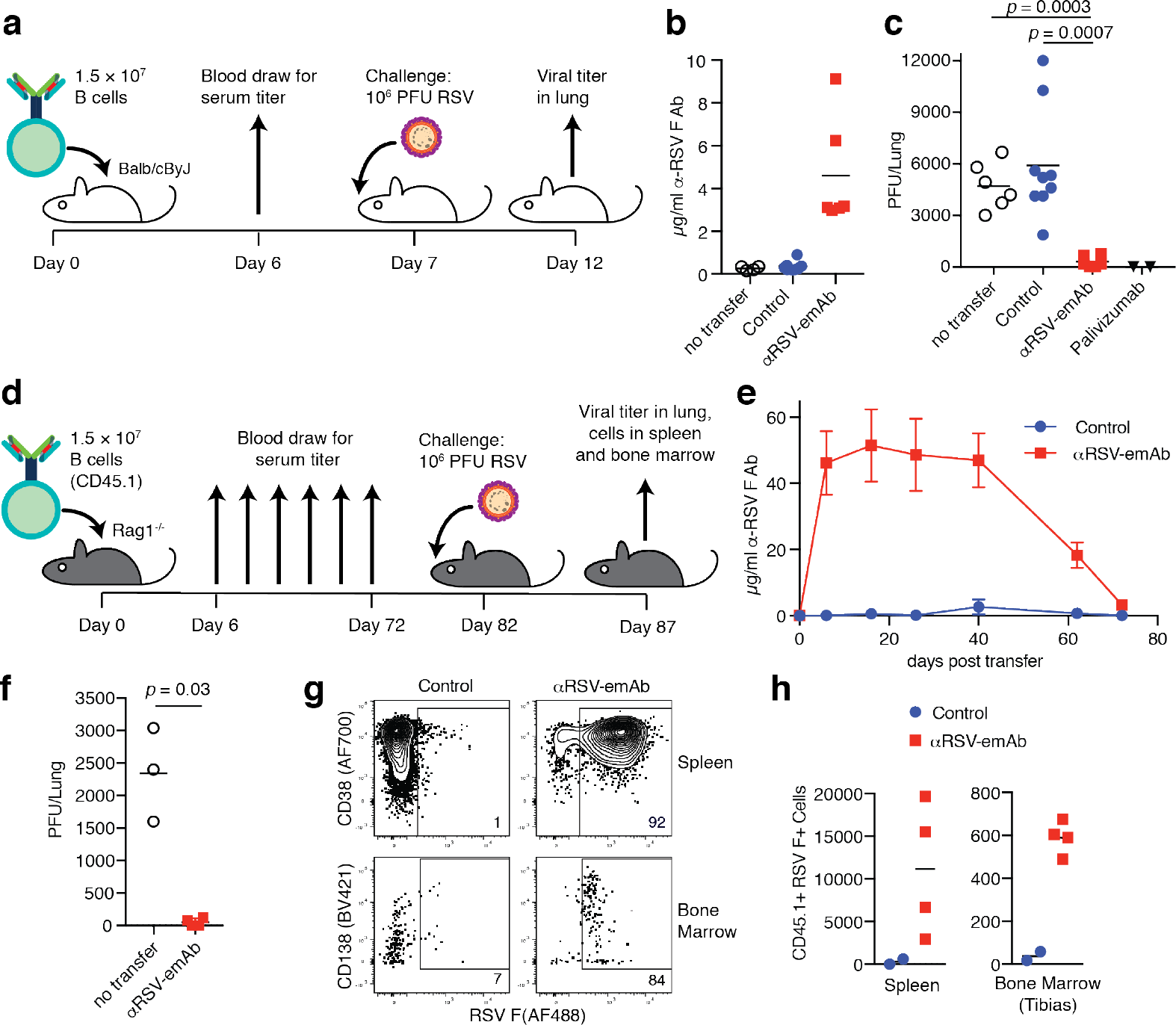
In vivo protection from RSV infection by αRSV-emAb B cells. (a) Schematic representation of experiment to test acute antiviral protection by αRSV-emAb B cells seven days following transfer into Balb/cByJ recipient mice. (b) ELISA-mediated quantitation of RSV F antigen-specific antibodies in serum from control mice that did not receive transfer, or from mice six days after the transfer of control B cells or αRSV-emAb B cells. (c) RSV titers in the lungs of mice who received no B cell transfer, control B cells, αRSV-emAb B cells, or 15 mg/kg palivizumab followed by intranasal infection with 10^6^ PFU of RSV (n=2-9). (d) Schematic representation of experiment to test long-term antiviral protection by transferred αRSV-emAb B cells in Rag1^−/−^ mice. (e) RSV F antigen-specific antibodies in serum in Rag1^−/−^ mice after the transfer of control or αRSV-emAb B cells (n=2-4). (f) RSV titers in the lungs of control mice who received no cell transfer compared to mice that received αRSV-emAb B cells 82 days prior to infection. (g) Representative flow cytometric analysis of CD38 and CD138 expression and (h) quantitation of transferred CD45.1^+^ control or αRSV-emAb B cells from spleen and bone marrow in mice 5 days after infection, 87 days after cell transfer.

One of the most susceptible groups for RSV infection are hematopoietic stem cell transplant recipients. Due to the profound immunodeficiency resulting from transplant, RSV infection in the three month period post-transplant carries a significant risk of lower respiratory tract infection, pneumonia, and death^31^. For this reason, we tested the capacity of αRSV-emAb B cells to provide long-term protection in the immunodeficient *Rag1*^−/−^ mice, which lack B and T cells (Fig. 6d). Transfer of 1.5 × 10^7^ CD45.1^+^ C57Bl/6 αRSV-emAb B cells into *Rag1*^−/−^ mice led to a rapid accumulation of sustained titer of above 40 μg/mL RSV-specific antibodies, which remained at this level for 40 days (Fig. 6e). Beginning at day 40, titers declined to ~5 μg/mL 72 days after cell transfer (Fig. 6e). Despite this decline in titers, intranasal challenge of mice with RSV 82 days after αRSV-emAb B cell transfer revealed near complete protection from infection (Fig. 6f). Analysis of transferred CD45.1^+^ cells after RSV infection revealed thousands of CD38^+^ memory αRSV-emAb B cells in the spleen, and hundreds of CD138^+^ αRSV-emAb-secreting cells in the bone marrow (Fig. 6g, h, Extended Data Fig. 1). Together, these results demonstrate that B cells can be efficiently engineered to provide robust and durable protection against infection.

## Discussion

The isolation of monoclonal antibodies has transformed medicine as therapeutics^32–34^. However, the use of antibody-producing primary B cells in adoptive cellular therapy has lagged behind that of other cell types. There has been some recent genetic engineering work focused on taking advantage of the potent protein secretion capabilities of B cells to produce non-antibody therapeutic proteins^35,36^, but the immense sequence diversity of the antibody repertoire has represented a major barrier to genome engineering of B cell specificity. We bypassed the sequence diversity obstacle by targeting an intronic region, allowing for universal B cell engineering without *a priori* knowledge of the endogenous *VDJ* sequences. Insertion of emAb cassettes into this region was complicated by the ability of B cells to simultaneously express productive heavy chains from both alleles, which allowed the emAb to be expressed from one allele and an endogenous heavy chain from the other. Endogenous antibody/emAb co-expression can be circumvented by selecting for pure emAb-expressing B cells by selecting against B cells that express both kappa and lambda light chains when the endogenous antibody and emAb antibodies express one or the other. Alternatively, B cells can be engineered and selected based upon the simultaneous expression of two emAbs. The latter strategy is intriguing since complimentary epitopes on the same pathogen or escape mutations could be simultaneously targeted by the same engineered cells.

Engineered emAb receptors are not limited by the gene segments and recombination events that generate the endogenous B cell repertoire. This could be important in situations where current vaccines have failed. For example, many of the broadly neutralizing antibodies which have been identified for HIV-1 contain features which are rare in the naive B cell repertoire^37^. As an alternative to isolation of rare antibodies, targeting domains could be designed *in silico* based either on the backbone of antibody variable domains, or entirely novel high affinity binding domains^38,39^. Combining these approaches with the emAb platform could allow targeting of pathogens for which no protective antibody has been isolated.

For high risk populations such as those with immunodeficiencies, common viral infection frequently leads to hospitalization, disability, and death. Hematopoietic stem cell recipients are a particularly relevant group since they are vulnerable to infection post-transplant and are already receiving a cellular product as part of treatment for an underlying disease. If some donor B cells were engineered and infused as emAb B cells targeting RSV, HMPV, EBV and CMV, thousands of hospital visits, disabilities, and deaths could be prevented each year. For lower risk populations, *in vitro* culture and infusion of engineered cells could pose a barrier to the clinical translation of emAb B cells. However, new technology is being developed to bypass patient-specific *in vitro* preparation of adoptive cellular therapies such as the production of universal donor cells^40^, as well as nanocarrier driven *in vivo* transduction of primary cells^41^.

In summary, we have demonstrated specific and efficient engineering of primary mouse and human cells to produce multiple potent antiviral antibodies using the emAb platform. Modified heavy chain loci in these engineered B cells retain the ability to undergo alternative splicing to produce both cell surface BCR and secreted antibodies and produced protective levels of antibody following infusion into mice. This technique offers the possibility of engineering humoral immunity to produce sterilizing immunity to diseases for which no current therapy exists.

## Acknowledgments

We thank M. D. Grey and L. Stamatatos for providing HIV-1 Env, plasmids containing the VRC01 sequence and purified VRC01 antibody, A. T. McGuire for EBV G_H_/G_L_, plasmids containing AMM01 and purified AMM01 antibody., B. Graham for plasmids encoding RSF F prefusion antigen, P. Kwong for plasmids encoding influenza HA stem, D. Stone for help producing AAVs, P. Collins and U. Buchholz for RSV expressing eGFP, M. J. McElrath for PBMCs from Seattle Area Control cohort, and K. G. Anderson for critical reading of the manuscript. Research reported in this publication was supported by an Individual Biomedical Research Award from The Hartwell Foundation (J.J.T.), a Sponsored Research Agreement from Vir Biotechnology (J.J.T.), and the National Institutes of Health under award numbers T32AI118690 (J. B.) and T32GM095421 (K.S.F.). The content is solely the responsibility of the authors and does not necessarily represent the official views of the National Institutes of Health.

## Author contributions

H.F.M. and J.J.T. designed the study, analyzed results, and wrote the manuscript with help from C.H. and J. B.; H.F.M., C.H., and K.S.F. performed and analyzed electroporations, adoptive transfers, and flow cytometry experiments; J.B. performed and analyzed plaque assays and produced and purified RSV F antigen and infectious RSV; M.T., C.H. and H.F.M. produced and purified AAVs.

## Methods

### Cell lines

3T3-msCD40L were obtained from Dr. Mark Connors at the NIH AIDS Reagent Program, Division of AIDS, NIAID, NIH (Cat#12535) and cultured in DMEM medium with 10% fetal calf serum (Gibco), 100 U/ml penicillin plus 100 μg/mL streptomycin (Gibco), and G418 (350 μg/mL). RAMOS cells were obtained from ATCC (CRL-1596™) and cultured in RPMI medium with 10% fetal calf serum (Gibco) and 100 U/mL penicillin plus 100 μg/mL streptomycin (Gibco). HEK293 cells were obtained from ATCC (CRL-1573™) cultured in DMEM medium with 10% fetal calf serum (Gibco), 100 U/mL penicillin plus 100 μg/mL streptomycin (Gibco). Vero cells were obtained from ATCC (CCL-8™) and cultured in DMEM medium with 10% fetal calf serum (Gibco), 100 U/mL penicillin plus 100 μg/mL streptomycin (Gibco).

### Design of single-chain antibody templates sequences

Human emAb cassettes consisted of a 450 base pair homology arm corresponding to 105,862,885-105,863,334 of human chromosome 14, the *IgVH1-69* heavy chain promoter region 106,715,182-106,715,481, the full-length antibody light chain gene, a segment encoding a 57 amino acid glycine-serine linker containing 3 tandem copies of the Streptag-II motif, the variable region of the heavy chain, and a splice junction with 60 base pairs of flanking sequence derived from matching *IgHJ* variable regions followed by a 450 base pair homology arm corresponding to 105,862,435-105,862,884 of human chromosome 14. Antibody variable domain sequences were derived from the humanized monoclonal antibody MEDI-493/palivizumab^6^, and the human monoclonal antibodies AMM01, VRC01, and MEDI8852^26–28^.

Murine emAb cassettes consisted of an upstream homology arm, the *J5558H10* heavy chain promoter region, 114,060,274 to 114,060,592 of chromosome 12^19^, full length codon optimized antibody light chain, a segment encoding a 57 amino acid glycine-serine linker containing three tandem copies of the Streptag-II sequence, codon optimized variable region of the heavy antibody chain, and a splice junction with 60 base pairs of flanking sequence derived from the mouse *IGHJ3* gene segment followed by a downstream homology arm. Antibody variable domains were derived from mouse Mab 1129^6^. The AAV emAb cassette included a 503 base pair upstream homology arm and a 968 base pair downstream homology arm that corresponded to 113,428,131 to 113,428,632 and 113,427,163 to 113,428,130 of murine chromosome 12 respectively. The dsDNA emAb cassette included a 36 base pair upstream and downstream homology arms that corresponded to 113,428,128 to 113,428,166 and 113,428,095 to 113,428,133 of murine chromosome 14, respectively.

### Production of recombinant emAb-AAV

To generate AAV plasmids for homologous recombination, linearized AAV backbone was isolated from pAAV-GFP (Addgene, Plasmid #32395) by digestion with SnaB (New England Biolabs), and homology arms for mouse and human heavy chain flanking an EcoRV restriction site inserted using NEB builder HiFi DNA assembly master mix (New England Biolabs). EmAb constructs were then synthesized as gene fragments (Integrated DNA Technologies) and cloned into the EcoRV site using Gibson HiFi master mix.

AAVs were generated by triple transfection of AAV emAb plasmid, serotype 6 capsid, and adenoviral helper plasmids into HEK293 cells using polyethylenimine (Polysciences). Eighteen hours after transfection, the media was changed to serum-free DMEM and the cells incubated for 48 hours cells prior to being lysed by freeze-thaw, treated with 20U of benzonase (Thermo Fisher Scientific) per 1 mL of viral lysate for 30 minutes at 37o C, then purified over iodixanol gradient. Purified AAV was concentrated into 1x DPBS using an Amicon Ultra-15 column (EMD Millipore)^42^ prior to viral titer determination by qPCR of AAV genomes^43^, which ranged from 5 × 10^10^ to 10^12^ per microliter.

### Production of murine double stranded DNA emAb templates

dsDNA templates containing short homology regions were generated from αRSV-emAb AAV plasmids through PCR amplification using Platinum PCR Supermix High Fidelity (Thermo Fisher Scientific) and modified DNA oligos. PCR product was purified and concentrated using minElute PCR cleanup columns (Qiagen). The following primers were used for amplification, with mouse genomic homology region in **bold** and phosphorothioate stabilized DNA bonds denoted by *:

Forward primer:

5’Phosphate/**ACCACCTCTGTGACAGCATTTATACAGTATCCGATG**GACAAGTGAGTGTCTCAGGTT AGGATTCT Reverse primer:

**T*A*A*AGAAAGTGCCCCACTCCACTCTTTGTCCCTATGC**TTGACCACAATGAATACTCCCACC

### Mouse B cell culture and electroporation

Mouse B cell medium consisted of RPMI supplemented with 10% Fetal calf serum (Gemini Biosciences), 10mM HEPES (Gibco), 55 μM Beta-mercaptoethanol (Sigma), and 100 U/ml penicillin plus 100 μg/mL streptomycin (Gibco) except in antibiotic free steps as noted. B cells were isolated from spleen and lymph nodes via negative selection with magnetic beads (Miltenyi Biotec) and 2×10^6^ cells/mL were cultured for 24 hours at 37oC in a tissue culture incubator in B cell medium supplemented with 100 ng/mL recombinant carrier free HA-tagged mouse CD40L (R&D systems), 100 ng/mL αHA antibody (clone 543851, R&D systems), and 4 ng/mL mouse IL-4 (R&D systems). Next, the B cells were electroporated using the Neon transfection system as follows: Cas9 protein (Invitrogen) and synthetic gRNA (Synthego) were precomplexed at a 1 to 3 molar ratio in Neon Buffer T at room temperature for at least 10 minutes. The muIgH_367_ gRNA sequence with the PAM site in bold is TTATACAGTATCCGATGCAT**AGG**. B cells were washed with 1xDPBS and suspended in Neon Buffer T at a final density of 2.5×10^7^ cells/mL with 12 μg of Cas9 per 10^6^ cells. When dsDNA emAb cassettes were used, 7.5 μg dsDNA template per 10^6^ cells was included in the electroporation. Cells were electroporated with three 10 millisecond pulses at 1675 volts and immediately dispensed into pre-warmed antibiotic-free mouse B cell medium. For AAV experiments, concentrated AAV in 1xDPBS was added up to 20% of final culture volume for 1 hour. After electroporation, B cells were expanded for an additional 48 hours with B cell medium supplemented with 100 ng/mL recombinant carrier free HA-tagged mouse CD40L, 100 ng/mL αHA antibody, 4 ng/mL mouse IL-4 (R&D systems), and 20 ng/mL mouse IL-21. (Biolegend). For additional expansion, B cells were co-cultured with irradiated (80 gy) NIH 3T3-CD40L feeder cells in the presence of 20 ng/mL mouse IL-21 for 6-8 days, with passage onto fresh irradiated 3T3-CD40L feeder cells every 4 days.

### Human B cell culture and electroporation

Human B cell medium was IMDM supplemented with 10% FBS (Gemini Biosciences), 100 U/mL penicillin and 100 μg/mL streptomycin (Gibco), except in antibiotic free steps as noted. Blood was obtained from healthy, HIV-seronegative adult volunteers as a part of the General Quality Control study in Seattle, WA by venipuncture and was approved by the Fred Hutch Institutional Review Board.

Informed consent was obtained before enrollment. PBMCs were isolated from whole blood using Accuspin System Histopaque-1077 (Sigma-Aldrich) resuspended in 10% Dimethylsulfoxide in heat-inactivated Fetal Bovine Serum and cryopreserved in liquid nitrogen before use. PBMCs were thawed and B cells isolated using negative selection using the Human B Cell Isolation Kit II (Miltenyi Biotec) according to the manufacturer’s recommendations. Isolated B cells were resuspended at 0.5-1.0×10^6^ cells/mL in stimulation media, which consisted of human B cell medium supplemented with 100 ng/mL MEGACD40L (Enzo Life Sciences), 50 ng/mL recombinant IL-2 (Biolegend), 50 ng/mL IL-10 (Shenandoah Biotech), 10 ng/mL IL-15 (Shenandoah Biotech), 1 μg/mL CpG ODN 2006 (IDT). After 48 hours, cells were electroporated using the Neon Transfection System. Cas9 protein (Invitrogen) and gRNA (Synthego) were precomplexed at a 2 to 1 molar ratio in Neon Buffer T for 20 minutes at room temperature. Cells were washed with 1xDPBS and resuspended to 2.5×10^7^ cells/mL in Neon Buffer T containing 12 μg of pre-complexed gRNA/Cas9 per 106 cells. The huIgH_296_ gRNA sequence with the PAM site in bold is GTCTCAGGAGCGGTGTCTGT**AGG**. The cell/gRNA/Cas9 mixture was electroporated with one 20 millisecond pulse at 1750V and immediately plated into stimulation media as described above, without antibiotics. After 30 minutes, AAV was added to a final concentration of up to 20% culture volume and mixed thoroughly. After 2-4 hours, cells were transferred to a larger culture dish to allow for further expansion. Two days after electroporation, cells were labeled with fluorochrome labeled antigen and/or streptactin and engineered cells FACS-purified. For secondary expansion, B cells were co-cultured for 6-12 days with irradiated (80 gy) NIH 3T3-CD40L feeder cells in human B cell medium containing 5 μg/mL Human recombinant Insulin (Sigma), 50 μg/mL Transferrin (Sigma), 50 ng/mL human IL-2 (Biolegend), 20 ng/mL human IL-21 (Biolegend), and 10 ng/mL human IL-15 (Shenandoah Biotech). Cells were passaged onto fresh 3T3-CD40L feeder cells every 4 days. In order to promote differentiation to plasma cells, cells were washed, and transferred from expansion conditions into fresh feeder-free culture conditions containing human B cell medium supplemented with 5 μg /mL Human recombinant Insulin (Sigma), 50 μg/mL Transferrin (Sigma), 500 U/mL Universal Type I IFN Protein (R&D Systems), 50 ng/mL IL-6 (Shenandoah Biotech), and 10 ng/mL IL-15 (Shenandoah Biotech).

### Assessment of gRNA activity by Sanger sequencing

Total genomic DNA was isolated from 0.5-2 × 10^6^ mock and cas9/gRNA treated cells two-five days following electroporation using the DNeasy kit (Qiagen). The genomic DNA region flanking the gRNA target site was amplified by PCR using the following primers:

Mouse IgH Forward, GGCTCCACCAGACCTCTCTA
Mouse IgH Reverse, AACCTCAGTCACCGTCTCCT
Human IgH Forward, ACAGTAAGCATGCCTCCTAAG
Human IgH Reverse, GCCACTCTAGGGCCTTTGTT

The resulting PCR product was purified using minElute reaction cleanup kit (Qiagen) and Sanger sequenced (Genewiz). The frequency of indels in Cas9/gRNA electroporated cells relative to mock electroporated cells was determined using the ICE algorithm^44^.

#### Protein antigens

Pre-fusion RSV F antigen trimer, EBV gH/gL complex, and a modified HIV Envelope GP140 trimer (426c TM4ΔV1-3) were produced as described^28,45,46^. Stabilized influenza HA stem was produced from VRC clone 3925, derived from strain H1 1999 NC as described^47^. All antigens were conjugated to Biotin NHS ester (Thermo Fisher Scientific) followed by tetramerization with streptavidin-PE, streptavidin-APC or streptavidin (all from Prozyme) as described previously^48^. RSV F antigen was conjugated to Alexa Fluor 488 NHS ester (Thermo Fisher Scientific) according to the manufacturer’s recommendations and used for flow cytometry.

### Flow Cytometry

Cells were incubated in 50 μL of FACS buffer containing a cocktail of antibodies for 30 minutes on ice prior to washing and analysis on a FACSymphony (BD Bioscience) or sorted on FACS Aria II (BD Bioscience). FACS buffer consisted of DPBS containing 5 mM EDTA and either 1% newborn calf serum (Life Technologies) or 1% bovine serum albumin (Sigma) for experiments including streptactin staining. For murine experiments, cells were labeled with a cocktails including a combinations of streptactin (IBA Lifesciences), anti-CD45.1 (A20, Biolegend), anti-CD138 (281-2, Biolegend), anti-B220 (RA3-6B2, BD Biosciences), anti-CD38 (90, eBioscience), GL7 (GL7, BD Biosciences), anti-CD19 (ID3, BD Biosciences), anti-CD3e (145-2C11, BD Biosciences), anti-F4/80 (BM8, Biolegend) and a fixable viability dye (eBioscience) prior to analysis. For human experiments, cells were labeled with a cocktail including a combinations of streptactin, anti-IgL-λ (MHL-38, Biolegend), anti-IgL-κ (G20-193, BD Biosciences), anti-CD19 (GSJ25C1, BD Biosciences), anti-CD20 (2H7, BD Biosciences), anti-CD27 (LG.7F9, eBioscience), anti-CD38 (HB-7, Biolegend), anti-CD45, (HI30, BD Biosciences), anti-CD69 (H1.2F3, Biolegend), anti-CD138 (MI15, BD Biosciences) and a fixable viability dye.

For *ex vivo* analysis of transferred cells after RSV infection in mice, single cell suspensions of spleen and bone marrow were generated by manual disassociation and filtration. Cells were stained with anti-CD45.1 APC (A20, Biolegend) and purified anti-CD16/32 (2.4G2, BioXcell) for 30 minutes on ice, washed with FACS buffer and then incubated with 25 μL of anti-APC conjugated magnetic microbeads (Miltenyi Biotec). Following a 15 - 30 minute incubation on ice, 3 mL of FACS buffer was added and the sample was passed over a magnetized LS column (Miltenyi Biotec). The tube and column were washed once with 5 mL of FACS buffer and then removed from the magnetic field. Five mL of FACS buffer was pushed through the column with a plunger twice to elute column-bound cells. Cells from the column-bound and 1/40 of the column flow through fractions were stained as described above. 20,000 AccuCheck counting beads (Invitrogen) were added to the samples to calculate total cell numbers.

Calcium flux was measured by flow cytometry with the Fluo-4 Direct kit (Thermo Fisher Scientific). Briefly, 1.5×10^6^ cells were labelled with 1 mL Fluo-4 according to manufacturer instructions for 30 minutes at 37oC. Fluo-4 baseline florescence was measured for 60 seconds, then cells were stimulated with 1 μg tetramerized RSV F antigen, followed by 180 seconds of measurement, and finally cells were simulated with 1 μg ionomycin in DMSO, and florescence was measured for an additional 60 seconds. Florescence intensity data was binned by time and displayed as fold change over baseline measurement. Flow cytometry data was analyzed using FlowJo × software (Tree Star).

### Animals

Animal studies were approved and conducted in accordance with the Fred Hutchinson Cancer Center Institutional Animal Care and Use Committee. Six- to ten-week old male and female BALB/cByJ, BALB/cByJ-CD45.1 congenic, C57bl/6-Rag1−/− (*Rag1*^−/−^), and C57bl/6-CD45.1 congenic mice were obtained from the Jackson Laboratory. For transfer of emAb B cells, age matched BALB/cByJ mice or *Rag1*^−/−^ mice received a single intraperitoneal (IP) injection of emAb or control B cells derived from CD45.1^+^ congenic donor mice, or a single IP injection of 15 mg/kg palivizumab at the indicated two days prior to RSV challenge.

### RSV infections and titer measurement

In RSV challenge experiments, mice were inoculated intranasally with 10^6^ PFU of sucrose purified RSV expressing eGFP^49^ in 40 μL 1xDPBS. Lungs were harvested five days post-infection and the titer was determined using a plaque assay. In brief, lungs were homogenized in 2 mL of DMEM using a GentleMACS M dissociator using preset program lung_02 (Miltenyi Biotec) followed by centrifugation at 400xg for 10 minutes. Supernatant was flash frozen and stored at −80°C. The supernatant was diluted 1:10 and 1:20 in DMEM and 100 μL of each dilution was added duplicate to confluent Vero cells in 24 well flat-bottoms tissue-culture plates and incubated for 2 hours at 37°C. An overlay of 0.8% methylcellulose was then added and plates incubated for five days prior to imaging on a Typhoon imager (GE Healthcare) with filter settings for eGFP. The titer in pfu/lung was calculated by counting the number of eGFP^+^ plaques with ImageJ software in the highest positive dilution and correcting for the dilution factor.

### ELISA

Nunc maxsorp 96-well plates (Thermo Fisher) were coated with 1 μg/mL of RSV F antigen, HIV Envelope, or influenza HA stem in 1xDPBS overnight at 4oC. Plates were washed three times with 1×DPBS containing 0.1% TWEEN-20 (PBST), and blocked with150 uL/well PBST+3% bovine serum albumin (BSA) (Sigma-Aldrich) for 1 hour at room temperature. Alternatively, 1 μg/mL of EBV gH/gL in 1x DPBS was coated on pre-blocked 96-well Ni-NTA plates (Qiagen) for 1 hour at room temperature, then washed three times with PBST. Antigen coated plates were incubated with culture supernatant or mouse plasma samples in duplicate diluted in PBST+3%BSA and a standard curve generated using the purified recombinant mouse RSV-specific antibody Mab 1127, or purified recombinant human RSV-specific palivizumab (Synagis clinical grade, MedImmune), Flu HA stem-specific Medi8852, EBV gH/gL-specific AMM01, or HIV-1 Envelope-specific VRC01 positive control antibodies for 90 minutes at room temperature. Plates were washed five times with PBST prior to 1 hour incubation with Horseradish peroxidase-conjugated goat anti-mouse or anti-human total Ig (Southern Biotech) diluted 1:4000 in PBST+3%BSA. Plates were then washed three times with PBST prior to a 2-15 minute incubation with 100 uL/well of ELISA TMB substrate (Thermo Fisher) and absorbance was measured at 405 nM using a Softmax Pro plate reader (Molecular Devices). The concentration of antigen-specific antibody in each sample was determined by reference to the standard curve and dilution factor.

### Statistical Analysis

Statistical analysis was performed using GraphPad Prism 7. Pairwise statistical comparisons were performed using unpaired two-tailed t-test with Welch’s correction. Unless otherwise indicated, values are given as mean and the error bars represent standard deviation.

---

**Extended Data Figure 1.**
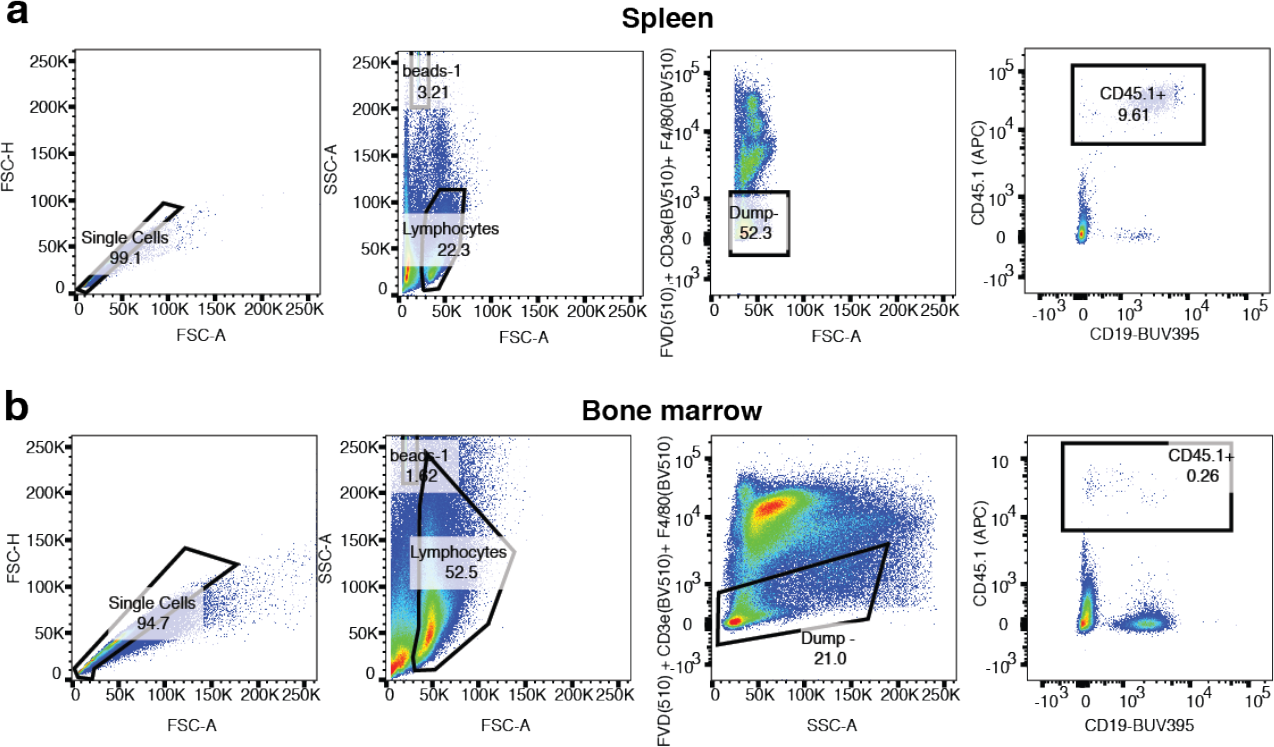
Identification of transferred CD45.1^+^ cells in spleen and bone marrow. Schematic showing exemplary nested gating of cells from spleen (**a**) and bone marrow (**b)**on single cells, lymphocytes, Dump channel (FVD-510, CD3e-BV510, and F4/80-BV510) negative, and CD45.1 positive cells. Position of AccuCheck beads is shown on the lymphocyte gate.

